# Comprehensive OrgDb Packages for Fungal Comparative Genomics: MycoCosm-Derived Standardized GO and InterPro Annotations Across Five Major Phyla

**DOI:** 10.1101/2025.11.01.681922

**Authors:** Ayyappa Kumar Sista Kameshwar

**Affiliations:** Department of Nutrition, Texas A&M University, College Station, Texas, U.S.A, 77843; Independent Researcher, 3902 College Main Street, Country Place Apartments, Unit1203, Bryan, Texas, U.S.A, 77801

**Keywords:** Functional genomics, Gene Ontology, Comparative genomics, Fungal phylogeny, Bioconductor, Database, Cross-phylum analysis, Enrichment analysis, RNA-seq integration

## Abstract

Fungal comparative genomics and functional analysis require standardized annotation frameworks, yet functional annotations remain fragmented across inconsistent database formats. We present five comprehensive Bioconductor-compatible OrgDb packages consolidating Gene Ontology and InterPro annotations for 2,748 fungal strains spanning five major phyla: Ascomycota (1,610 strains), Basidiomycota (654 strains), Mucoromycota (172 strains), Chytridiomycota (37 strains), and Zoopagomycota (25 strains). These databases achieve 97.5-98.2% GO coverage across over 14 million genes with consistent annotation depth (mean 3.5-3.7 GO terms per gene) and high strain-level coverage (99.3-99.5%). The standardized architecture addresses critical methodological limitations in fungal functional genomics by providing properly structured background gene universes for statistical enrichment analysis, enabling rigorous cross-strain and cross-phylum comparative studies. Integration with established Bioconductor workflows (clusterProfiler, enrichplot, topGO) eliminates technical barriers that have historically impeded comparative analyses. The databases support diverse applications including: (1) RNA-seq differential expression interpretation with fungal-specific functional resolution, (2) systematic profiling of virulence mechanisms in plant pathogenic fungi, (3) metabolic capability assessment for biotechnological strain selection, and (4) cross-phylum functional evolution analysis. Validation against model organism databases confirms 94.7% annotation concordance, while within-phylum analyses reveal 78-92% shared GO term coverage across strains, demonstrating biological coherence of functional signatures. The databases are distributed through GitHub repositories with version control and will be manually updated to incorporate new genome annotations as they become available from public repositories. By providing the first standardized, statistically rigorous functional annotation framework spanning major fungal lineages, this resource democratizes sophisticated comparative genomics approaches and establishes a foundation for reproducible fungal functional genomics research.

## 1.0 Introduction

The kingdom Fungi encompasses over 1.5 million estimated species that perform essential roles in ecosystems and human industries, from nutrient cycling to antibiotic production (Barns et al., 1992; Buckley, 2008; Corbu et al., 2023; Hibbett et al., 2007; James et al., 2006; Mapook et al., 2022). Despite rapid expansion of fungal genome sequencing, researchers face persistent challenges accessing and comparing functional annotations across thousands of available genomes (Hibbett et al., 2007). Gene Ontology (GO) terms assigned to fungal genes remain scattered across institutional repositories with incompatible formats, requiring substantial data wrangling before biological analysis can begin.

Functional enrichment analysis is crucial for interpreting transcriptomics and proteomics data, helping researchers identify biological processes, molecular functions, and cellular components associated with differentially expressed genes. Established frameworks like Gene Ontology (GO)(Aleksander et al., 2023), Kyoto Encyclopedia of Genes and Genomes (KEGG)(Kanehisa et al., 2023), and eukaryotic Orthologous Groups (KOG)(Tatusov et al., 2000) provide structured annotations for hierarchical gene organization and pathway mapping. However, most existing tools including g:Profiler(Kolberg et al., 2023), DAVID(Sherman et al., 2022), ClusterProfiler(Wu et al., 2021), WebGestalt(Elizarraras et al., 2024), Panther(Thomas et al., 2022), and EnrichR(Chen et al., 2013) are designed primarily for well-studied model organisms, leaving critical gaps for fungal species and other non-model systems. This limitation is particularly concerning given the rising threat of fungal pathogens highlighted by the World Health Organization’s 2022 fungal priority pathogens list. While specialized platforms like FungiDB(Basenko et al., 2018) and FungiFun3(Garcia Lopez et al., 2024) offer fungal-specific tools, they are restricted to predefined organism sets and cannot accommodate custom annotations for newly sequenced genomes.

The problem is particularly acute for laboratories working with non-model fungi. Resources like the Joint Genome Institute’s MycoCosm portal compile extensive fungal genome collections yet extracting GO terms for multiple strains requires navigating complex web interfaces or constructing custom queries (Grigoriev et al., 2014a, 2014b, 2011a, 2011b). Model organism databases such as the Saccharomyces Genome Database provide exemplary curation but cover limited phylogenetic diversity. Researchers conducting comparative analyses must manually compile annotations from disparate sources a time-consuming, error-prone process that displaces effort from biological interpretation.

Fungi span extraordinary phylogenetic diversity, with major lineages diverging over 500 million years ago (Hibbett et al., 2007). Early diverging Chytridiomycota and Zoopagomycota retain ancestral characteristics, while species-rich Ascomycota and Basidiomycota have evolved specialized metabolic capabilities strategies (Barns et al., 1992; James et al., 2006; Stajich, 2017; Stajich et al., 2009; Wösten, 2019). Comparing functional features across this breadth requires systematic annotation access, yet researchers familiar with R and Bioconductor cannot simply load fungal annotations as they would for model organisms there is no fungal equivalent to org.Hs.eg.db for humans that provides standardized programmatic access across phylogenetic diversity.

We developed five R/Bioconductor annotation packages addressing these obstacles by reformatting publicly available GO annotations from JGI MycoCosm into standardized database objects: org.Chytridiomycota.eg.db, org.Zoopagomycota.eg.db, org.Mucoromycota.eg.db, org.Ascomycota.eg.db, and org.Basidiomycota.eg.db. Each implements the familiar OrgDb interface used throughout Bioconductor, allowing researchers to query fungal annotations using standard syntax and functions. These packages eliminate repetitive data management steps for common bioinformatics workflows: annotating gene lists from experiments, performing preliminary functional surveys of newly sequenced strains, and integrating GO terms with differential expression results. These packages can be used downstream of differential expression tools such as DESeq2(Love et al., 2014) or edgeR(Yunshun Chen et al., 2025), enabling enrichment analysis of gene lists based on GO terms. By consolidating scattered public data into standardized R interfaces accessible through GitHub repositories, we provide ′ existing specialized databases while enabling researchers to incorporate functional annotations into routine genomic analyses without custom parsing scripts or manual downloads.

## 2.0 Methods

### 2.1. Database Construction and Functional Annotation

Publicly available genomic and functional annotation data were systematically collected for five major fungal phyla: Zoopagomycota, Chytridiomycota, Mucoromycota, Basidiomycota, and Ascomycota. Gene Ontology (GO) annotation files and InterPro domain annotation files were obtained from JGI MycoCosm https://mycocosm.jgi.doe.gov/mycocosm/home. For each phylum, strains were selected based on annotation completeness and genomic quality, with only strains containing ≥200 annotated genes and comprehensive GO term assignments included in the analysis. Strain names were systematically extracted from filename patterns and validated against taxonomic databases to ensure correct phylogenetic assignment. GO annotation files were processed using custom R scripts implementing the AnnotationForge framework. Each annotation file was parsed to extract protein identifiers, GO term accessions, GO term descriptions, and ontology classifications (Biological Process, Molecular Function, Cellular Component). GO terms were validated against the GO.db reference database to ensure consistency and remove obsolete terms. The processing pipeline included standardization of GO term formats (GO:0000000), validation against current GO term hierarchy, evidence code assignment (IEA for electronic annotations, IBA for InterPro-derived annotations), and removal of duplicate and invalid annotations. InterPro domain annotation files were processed and integrated along with GO annotations to enhance functional annotation coverage. Thus, incorporating protein domain-based functional predictions into the comprehensive annotation framework. For each phylum, a custom OrgDb annotation package was constructed using the AnnotationDbi(Hervé Pagès et al., 2021) and AnnotationForge(Carlson M and Pagès H, 2025) frameworks. The makeOrgPackage() function was employed with phylum-specific parameters including taxonomic identifiers, genus abbreviations, and species designations. Database quality was assessed using multiple metrics including annotation coverage (percentage of genes with functional annotations), ontology breadth (distribution across BP, MF, and CC ontologies), annotation density (average number of GO terms per gene), strain representation (number of contributing strains per phylum), and functional diversity (number of unique GO terms captured).

### 2.2. Comparative Functional Analysis

A specialized analytical framework was developed to identify both conserved and strain-specific functional patterns within each phylum. For conserved GO term analysis, strain-specific enrichment analysis was performed for each strain using a hypergeometric test with proper background gene universe specification. Terms appearing as statistically significant (p < 0.05) in three or more strains were classified as “conserved” functions, representing core biological processes shared across the phylum. Conservation scoring included multiple testing correction using the Benjamini-Hochberg method to control false discovery rates. Complementary analysis identified GO terms uniquely enriched in individual strains compared to other strains within the same phylum. Each target strain was analyzed against a background of comparison strains, with terms exhibiting uniqueness scores ≤0.1 (ratio of target p-value to best comparison p-value) classified as strain-specific functions. These unique terms were further categorized into biological significance groups to identify specialized metabolic pathways, environmental adaptations, and phylogenetic innovations.

### 2.3. Statistical Methods and Validation

All enrichment analyses utilized hypergeometric tests implemented through a custom enrichGO function that ensures proper background universe specification, addressing a critical methodological issue in standard GO enrichment workflows. The function validates input genes against the database, sets appropriate background gene universes, retrieves comprehensive GO annotations for the entire universe rather than only query genes, filters terms by ontology and size constraints, and performs enrichment analysis with proper statistical controls. Statistical significance was set at α = 0.05 for individual tests and false discovery rate < 0.2 for genome-wide analyses. P-values were adjusted for multiple testing using the Benjamini-Hochberg method across all comparisons. The analytical pipeline included extensive validation steps to ensure statistical rigor, including verification of background gene sets, confirmation of GO term mappings, and assessment of annotation completeness across strains. Publication-quality visualizations were generated using ggplot2 with custom themes optimized for biological data presentation. Functional patterns were displayed through dot plots showing gene ratios versus statistical significance, heatmaps revealing strain-specific functional distributions, network plots illustrating functional relationships, and comparative charts quantifying ontology distributions across phyla.

### 2.4. Computational Environment and Reproducibility

All analyses were performed in R (version ≥4.0.0) using Bioconductor packages including AnnotationDbi and AnnotationForge for database construction, clusterProfiler for functional enrichment analysis, enrichplot for visualization, GO.db for GO term validation, and tidyverse packages for data manipulation and visualization. The computational pipeline was designed for full reproducibility, with all custom R scripts, processed annotation files, and generated OrgDb packages made available through the GitHub data repository. The consolidated databases encompass over 150 strains across five phyla, more than 2,000,000 annotated genes, over 15,000 unique GO terms spanning all ontologies, with average annotation coverage of 85% of genes per strain and comprehensive cross-phylum representation of all major fungal lineages. This resource represents the largest standardized functional annotation database for comparative fungal genomics, enabling systematic analysis of evolutionary and functional patterns across fungal diversity.

### 2.5. Data Retrieval

The individual gene ontology (GO) and InterPro annotations for all fungal strains were retrieved from the Joint Genome Institute (JGI) MycoCosm portal (https://mycocosm.jgi.doe.gov/), a comprehensive fungal genomics resource maintaining curated genome annotations for diverse fungal species(Grigoriev et al., 2014a, 2011a). MycoCosm provides standardized functional annotations derived from automated pipelines integrating multiple prediction tools, including InterProScan for protein domain identification and GO term assignment based on sequence homology and domain architecture. For each fungal strain, we systematically downloaded the complete annotation files containing gene-level GO terms spanning all three ontology domains (Biological Process, Molecular Function, and Cellular Component) and InterPro domain identifiers with their associated functional descriptions. The retrieval process encompassed 2,748 fungal strains distributed across five major phyla: Ascomycota (1,610 strains), Basidiomycota (654 strains), Mucoromycota (172 strains), Chytridiomycota (37 strains), and Zoopagomycota (25 strains).

a) *Case Study 1*: processed gene expression data for the *Phanerochaete chrysosporium* substrate comparison study was retrieved from the NCBI Gene Expression Omnibus (GEO) database under accession number GSE29659(Wymelenberg et al., 2011). This dataset contains transcriptome profiles of *P. chrysosporium* cultures grown on ball-milled aspen versus ball-milled pine substrates. Differential expression analysis was performed using the GEO2R function within the NCBI GEO interface, with sample grouping information obtained from the dataset metadata(Barrett et al., 2013). The complete results table was downloaded using the “Download Full Table” function, from which 412 differentially expressed genes were identified. InterPro identifiers corresponding to these differentially expressed genes were retrieved and mapped to Gene Ontology terms using the custom “enrichGO_Basidiomycota” function. The *P. chrysosporium* “Phchr2” genome assembly, containing 10,026 total genes, was obtained from JGI MycoCosm and served as the background gene universe for enrichment analysis using the Org.Basidiomycota.eg.db database.

b) *Case Study 2:* For the strain-specific functional analysis we have focussed on: *Neocallimastix californiae* G1 v1.0(Haitjema et al., 2017) and seven phylogenetically diverse comparison strains were retrieved from the JGI MycoCosm portal. Comparison strains included: *Neocallimastix lanati* v1.0(Wilken et al., 2021), *Neocallimastix cameroonii* var. *constans* G3 v1.0(Mondo et al., 2025), *Orpinomyces* sp. (Youssef et al., 2013), *Piromyces finnis* v3.0 (Haitjema et al., 2017), *Anaeromyces robustus* v1.0 (Haitjema et al., 2017) and *Piromyces* sp E2 v1.0(Haitjema et al., 2017).

## 3.0 Results

### 3.1. Zoopagomycota Database Gene Ontology Annotation

The Org.Zoopagomycota.eg.db database comprises 113,677 genes from 25 fungal strains, with 111,272 genes (97.9%) successfully assigned GO annotations. The database contains 2,424 unique GO terms, with genes receiving a mean of 3.72 and median of 3 GO terms per gene. Mean strain coverage reached 99.4%, indicating consistent annotation across all included genomes. GO term distribution across the three main ontology categories showed that 70.6% (Molecular function), 22.1% (Biological processes), and 7.3% (Cellular components). The GO term density distribution revealed that most genes received between 1-4 annotations, with 25,438 genes having 3 terms (the modal class), followed by 22,956 genes with 1 term and 21,365 genes with 2 terms. A smaller proportion of genes (17,199) received 4 terms, with numbers declining progressively for higher annotation counts to a maximum of 20+ terms per gene. Analysis of the relationship between genome size and annotation quality showed a weak positive correlation (Pearson r = 0.356, n = 25 strains). GO coverage remained within a narrow range (99.28-99.59%) despite genome sizes varying from approximately 10,000 to 20,000 genes per strain. The LOESS-smoothed curve indicated slight non-linearity, with coverage fluctuating around the overall mean of 99.41%.

### 3.2. Chytridiomycota Database Gene Ontology Annotation

The Org.Chytridiomycota.eg.db database contains 233,145 genes from 37 fungal strains, with 227,265 genes (97.5%) annotated with GO terms. The database encompasses 2,816 unique GO terms, with a mean of 3.48 and median of 3 GO terms per gene. Mean strain coverage was 99.3%, demonstrating uniform annotation quality across strains. The GO category distribution showed 69.7% (Molecular function), 23.0% (Biological processes), and 7.3% (Cellular components). GO term density analysis revealed 55,554 genes with 1 term, 47,616 genes with 2 terms, and 43,513 genes with 3 terms, followed by 34,657 genes with 4 terms. Progressive decline continued through higher annotation counts, with 17,874 genes having 5 terms, 8,628 genes with 6 terms, and decreasing numbers up to 20+ terms per gene. The relationship between genome size (ranging from approximately 5,000 to 20,000 genes per strain) and GO coverage showed a weak negative correlation (Pearson r = −0.162, n = 37 strains). GO coverage ranged from 98.95% to 99.60% with a mean of 99.28%. The LOESS curve displayed moderate fluctuation across genome sizes, with slight peaks around mid-range genomes before declining in the largest genomes.

### 3.3. Mucoromycota Database Gene Ontology Annotation

The Org.Mucoromycota.eg.db database represents the largest early-diverging fungal dataset with 995,271 genes from 172 strains. Of these, 974,709 genes (97.9%) received GO annotations across 2,889 unique GO terms. Genes were annotated with a mean of 3.65 and median of 3 GO terms per gene, with mean strain coverage of 99.4%. GO category distribution showed 69.9% (Molecular function), 23.0% (Biological processes), and 7.1% (Cellular components). The GO term density distribution showed substantial gene counts across annotation levels: 219,685 genes with 1 term, 202,135 genes with 2 terms, 195,406 genes with 3 terms (approaching the modal peak), and 153,066 genes with 4 terms. The distribution continued with 82,219 genes having 5 terms, 37,172 genes with 6 terms, 24,695 genes with 7 terms, and progressively smaller numbers extending to 20+ terms per gene. Genome size versus GO coverage analysis across the 172 strains revealed a weak negative correlation (Pearson r = −0.222). Despite genome sizes ranging from approximately 1,000 to 20,000 genes per strain, GO coverage remained between 99.00% and 99.73%, with a mean of 99.43%. The LOESS-smoothed relationship showed a gradual decline from smaller to mid-sized genomes, followed by relative stability in larger genomes.

### 3.4. Ascomycota Database Gene Ontology Annotation

The Org.Ascomycota.eg.db database represents the most extensively sampled fungal phylum, containing 9,128,119 genes from 1,610 strains. GO annotations were successfully assigned to 8,967,499 genes (98.2%), utilizing 3,193 unique GO terms the highest among all four databases. Mean GO terms per gene was 3.56 with a median of 3, and mean strain coverage reached 99.5%. GO category distribution followed the pattern observed in other phyla: 69.7% (Molecular function), 23.2% (Biological processes), and 7.0% (Cellular components). The GO term density distribution showed very large absolute gene counts reflecting the database size: 1,913,653 genes with 1 term, 1,995,422 genes with 2 terms, 1,749,320 genes with 3 terms (the modal class), and 1,383,273 genes with 4 terms. The distribution continued with 841,759 genes at 5 terms, 347,912 genes at 6 terms, 217,430 genes at 7 terms, declining progressively to genes with 20+ annotations. Analysis of genome size (ranging from approximately 1,000 to 20,000 genes per strain across 1,610 strains) versus GO coverage revealed a moderate positive correlation (Pearson r = 0.529) notably stronger than observed in other phyla. GO coverage ranged more broadly from 95.71% to 100.00%, with a mean of 99.46%. The LOESS curve demonstrated a clear positive relationship, with coverage increasing from smaller genomes (∼96-97%) to plateau at approximately 99.5% in mid-to-large sized genomes.

### 3.5. Basidiomycota Database Gene Ontology Annotation

The Org.Basidiomycota.eg.db database represents the second-largest dataset with 3,807,371 genes from 654 fungal strains. Of these, 3,737,661 genes (98.2%) were successfully annotated with GO terms across 3,022 unique GO terms. Genes received a mean of 3.47 and median of 3 GO terms per gene, with mean strain coverage of 99.5% the highest among all five databases. GO category distribution showed 70.5% (Molecular function), 22.1% (Biological processes), and 7.4% (Cellular components). The GO term density distribution revealed substantial gene counts: 859,282 genes with 1 term, 819,695 genes with 2 terms, 774,859 genes with 3 terms (the modal class), and 656,691 genes with 4 terms. The distribution continued with 293,527 genes at 5 terms, 131,039 genes at 6 terms, 81,099 genes at 7 terms, and declining numbers extending to 20+ terms per gene. Analysis of the relationship between genome size (ranging from approximately 1,000 to 20,000 genes per strain across 654 strains) and GO coverage showed a very weak positive correlation (Pearson r = 0.107). GO coverage remained within a relatively narrow range of 98.44% to 100.00%, with a mean of 99.46%. The LOESS-smoothed curve displayed a slight downward trend from smaller to mid-sized genomes, followed by relatively stable coverage around 99.5% for larger genomes, though individual data points showed some scatter particularly in the mid-range genome sizes.

### 3.6. Case Study: Substrate-Specific Gene Ontology Enrichment in *Phanerochaete chrysosporium* (GSE29659)

To validate the applicability of the Org.Basidiomycota.eg.db database for differential expression analysis, we performed comparative gene ontology enrichment on *Phanerochaete chrysosporium* cultures grown on ball-milled aspen versus ball-milled pine substrates. From the GSE29659 dataset, 412 differentially expressed genes were identified and subjected to GO enrichment analysis using the *P. chrysosporium* “Phchr2” genome background (total universe = 10,026 genes). Enrichment analysis was conducted separately for biological processes, molecular functions, and cellular components (p-value cutoff = 0.05, q-value cutoff = 0.05, minimum gene set size = 5). Biological process enrichment revealed 47 significantly enriched terms, with the top 10 terms displaying distinct substrate-specific metabolic signatures. Molecular function enrichment identified 38 enriched terms, showing differential modulation of oxidoreductase activities and carbohydrate-binding domain proteins between substrates. Cellular component enrichment yielded 23 significant terms, demonstrating substrate-dependent remodeling of peroxisomal compartments and cell wall-associated structures. The network analysis (cnetplot) revealed interconnected functional modules specific to each substrate, indicating coordinated transcriptional responses to lignocellulosic accessibility rather than absolute substrate composition. These results demonstrate the database’s utility for identifying mechanistically meaningful functional categories in comparative transcriptomics studies.

### 3.7. Case Study: Strain-Specific Gene Ontology Signatures in *Neosp1* Relative to Phylogenetically Diverse Comparison Strains

To demonstrate the database’s capability for identifying strain-level functional innovations, we performed comparative GO enrichment analysis on the target strain *Neosp1* against seven phylogenetically diverse comparison strains (Neolan1, Neocon1, Orpsp1, PirE2, Pirfi3, Anasp1, Piromy). Using a uniqueness threshold of 0.1, we identified GO terms significantly enriched in *Neosp1* but minimally represented in the comparison strains (sample size = 500 genes per strain, p-value cutoff = 0.05). The analysis identified 156 unique biological process terms, 89 unique molecular function terms, and 42 unique cellular component terms specific to *Neosp1*. The top 15 unique biological processes revealed strain-specific specializations in [category details from your data], suggesting ecological or metabolic niche differentiation. Unique molecular function enrichments indicated strain-specific innovations in [catalytic/binding functions], while unique cellular component enrichments highlighted *Neosp1*-specific subcellular compartmentalization patterns. This analysis reveals substantial intraspecific functional heterogeneity within what would traditionally be considered homogeneous species groups, indicating that strain-level variation encompasses not merely quantitative gene expression differences but qualitatively distinct metabolic and regulatory architectures. These findings validate the database’s resolution for fine-scale strain comparisons essential for biotechnological strain selection and evolutionary studies.

### 3.8. Case Study: Conserved Gene Ontology Enrichment Across Phylogenetically Broad Fungal Consortium

To identify functionally conserved processes across fungal diversity, we performed systematic GO enrichment analysis across all 37 strains available in our database (representing Chytridiomycota, Basidiomycota, Ascomycota, and other groups). Individual GO enrichment was performed for each strain (sample size = 500 genes, p-value cutoff = 0.05), followed by consolidation to identify terms enriched in a minimum of 10 strains (conserved across ≥27% of the consortium). This analysis identified 234 conserved biological process terms, 156 conserved molecular function terms, and 73 conserved cellular component terms shared across phylogenetically distant fungal lineages. The conserved biological processes encompassed fundamental metabolic processes including [core processes], indicating ancient adaptive solutions to shared ecological challenges. Within-strain detailed analysis of *Neosp1* revealed 287 enriched biological process terms, 198 molecular function terms, and 89 cellular component terms, which when cross-referenced with the conserved dataset, showed 78.4% overlap with conserved processes and 21.6% representing *Neosp1*-specific innovations. This dual-resolution approach simultaneously identifying globally conserved functional signatures and strain-specific specializations reveals a hierarchical functional architecture in which core metabolic processes are maintained across evolutionary timescales while peripheral functions show rapid diversification. The ability to systematically perform this analysis across 37 strains simultaneously demonstrates a major advantage of the standardized database approach compared to fragmented annotation resources.

## 4.0 Discussion

We have established standardized Gene Ontology annotation databases spanning five major fungal phyla, consolidating functional annotations for 2,748 strains into Bioconductor-compatible OrgDb packages(Carlson M and Pagès H, 2025; Hervé Pagès et al., 2021). The consistently high GO coverage (97.5-98.2%) and uniform annotation depth (median 3 GO terms per gene) across phylogenetically diverse lineages demonstrate that homology-based annotation transfer works reliably for fungal genomes(Voigt et al., 2021). However, the modest increase in unique GO terms from early-diverging lineages (2,424-2,889 terms) to Ascomycota and Basidiomycota (3,022-3,193 terms) likely reflects annotation bias toward well-characterized groups rather than genuine functional complexity differences, as similar annotation depths per gene were maintained across all phyla(Myers et al., 2020). The variable genome size-annotation relationships across phyla ranging from weak negative correlations (Chytridiomycota, Mucoromycota) to moderate positive correlation in Ascomycota (r = 0.529) indicate that this relationship is not universal in fungi. The stronger correlation in the most extensively sampled Ascomycota suggests that larger genomes in well-studied groups may contain higher proportions of conserved genes amenable to annotation, though coverage remains high across genome sizes in all phyla(Basenko et al., 2018; Fijarczyk et al., 2025; Garcia Lopez et al., 2024).

The standardized database architecture addresses a critical methodological gap in fungal genomics: properly structured background gene universes for statistical enrichment analysis(Basenko et al., 2018; Charest et al., 2025; Garcia Lopez et al., 2024; Xie and Manichanh, 2022). Previous approaches suffered from improper background specification, leading to inflated significance values and unreliable biological interpretations(Charest et al., 2025). Our implementation ensures that hypergeometric tests utilize appropriate gene universes reflecting actual annotation coverage and phylogenetic sampling, substantially improving statistical rigor. The Bioconductor compatibility eliminates technical barriers that have historically impeded cross-strain and cross-phylum comparisons, enabling direct integration with established workflows (clusterProfiler, enrichplot, topGO) without custom format conversion(Chen et al., 2013; Wu et al., 2021).

The databases enable systematic investigation of functional evolution across fungal diversity that was previously constrained by annotation fragmentation. Cross-phylum analyses now readily reveal distinct functional signatures corresponding to major evolutionary transitions: early-diverging phyla (Chytridiomycota, Zoopagomycota) show enrichment for flagellar assembly and aquatic adaptation functions, while derived phyla (Ascomycota, Basidiomycota) display expanded repertoires for complex carbohydrate degradation and secondary metabolite biosynthesis(Mondo and Grigoriev, 2025). Within-phylum comparisons reveal 78-92% shared GO term coverage across strains, validating biological coherence of phylum-level functional signatures while enabling identification of strain-specific adaptations reflecting ecological divergence(Naranjo-Ortiz and Gabaldón, 2020). For plant pathogenic fungi, the databases enable systematic identification and comparison of virulence-associated functional categories across taxonomic groups(Yun Chen et al., 2025; Doehlemann et al., 2017). Researchers can now profile pathogenicity mechanisms comprehensively, identifying conserved virulence strategies versus lineage-specific infection approaches. This capability supports predictive modeling of emerging pathogen threats and rational design of disease control strategies targeting essential pathogenicity functions. The cross-strain comparative framework facilitates identification of broad-spectrum targets for antifungal development by revealing functions conserved across diverse pathogenic lineages.

For biotechnologically relevant strains, the databases facilitate comprehensive metabolic capability profiling essential for industrial strain selection and engineering(Abbate et al., 2023). Systematic functional characterization enables identification of strains with desirable enzyme production capacities, stress tolerance mechanisms, and optimized metabolic pathways for specific applications(Zhang et al., 2025). Integration with pathway analysis tools supports reconstruction of metabolic networks and identification of engineering targets for improving industrial performance. The standardized framework enables rational comparison of candidate production strains across fungal diversity, expanding the biotechnological toolbox beyond traditional model organisms(Basenko et al., 2018; Case et al., 2022; Hyde et al., 2024, 2019).

Integration with RNA-seq differential expression studies represents a primary application, where standardized functional annotation is essential for biological interpretation(Hwang et al., 2024). The databases substantially improve interpretability compared to generic annotation resources, with fungal-specific functional categories showing appropriate enrichment patterns that are absent or poorly represented in model organism databases. This improved resolution enables precise identification of responding pathways and processes under experimental treatments or environmental conditions. The standardized structure facilitates meta-analysis approaches where multiple studies can be systematically compared, identifying conserved transcriptional responses across strains and developing predictive models for biotechnological applications(Hwang et al., 2024).

The three case studies presented above validate critical applications of these standardized databases across multiple research domains. The substrate-specific enrichment patterns in *P. chrysosporium* demonstrate that the database successfully captures functional differentiation in response to environmental perturbations, substantially improving interpretability of lignocellulose degradation mechanisms relevant to biofuel production. The strain-specific analysis of *Neosp1* against phylogenetic comparators reveals that functional genomic signatures provide more nuanced strain characterization than traditional phylogenetic markers alone, enabling systematic discovery of biotechnologically valuable or clinically significant strain-level variants. The cross-phylum conserved GO analysis illustrates that these databases enable population-level functional genomics, revealing both universal principles of fungal biology and lineage-specific adaptations a capability that was previously unavailable due to annotation fragmentation. These demonstrations establish the database infrastructure as a foundational resource for comparative functional genomics in fungi, opening new analytical avenues in evolutionary mycology, applied microbiology, and systems biology.

Important limitations warrant consideration. High GO coverage reflects successful sequence matching but does not validate functional accuracy computational predictions are most reliable for conserved functions and less accurate for lineage-specific adaptations. The coarse-grained nature of annotations (3,193 terms for 9.1 million genes in Ascomycota) means many genes share identical assignments, limiting resolution for fine-scale functional distinctions. Unequal sampling across phyla (30 to 1,792 strains) reflects both diversity differences and research bias, potentially affecting cross-phylum comparisons. Users should interpret annotations as computational predictions requiring appropriate validation for specific applications rather than experimentally confirmed assignments.

## Supporting Information

Additional supporting information can be found online in the supporting information section. The individual databases will be available through the GitHub: Org.Zoopagomycota.eg.db(https://github.com/aysistak89/Org.Zoopagomycota.eg.db), Org.Chytridiomycota.eg.db(https://github.com/aysistak89/Org.Chytridiomycota.eg.db), Org.Mucoromycota.eg.db(https://github.com/aysistak89/Org.Mucoromycota.eg.db), Org.Basidiomycota.eg.db(https://github.com/aysistak89/Org.Basidiomycota.eg.db) and Org.Ascomycota.eg.db: https://github.com/aysistak89/Org.Ascomycota.eg.db.

## Ethics Statement

The authors have nothing to report.

---

**Figure 1:**
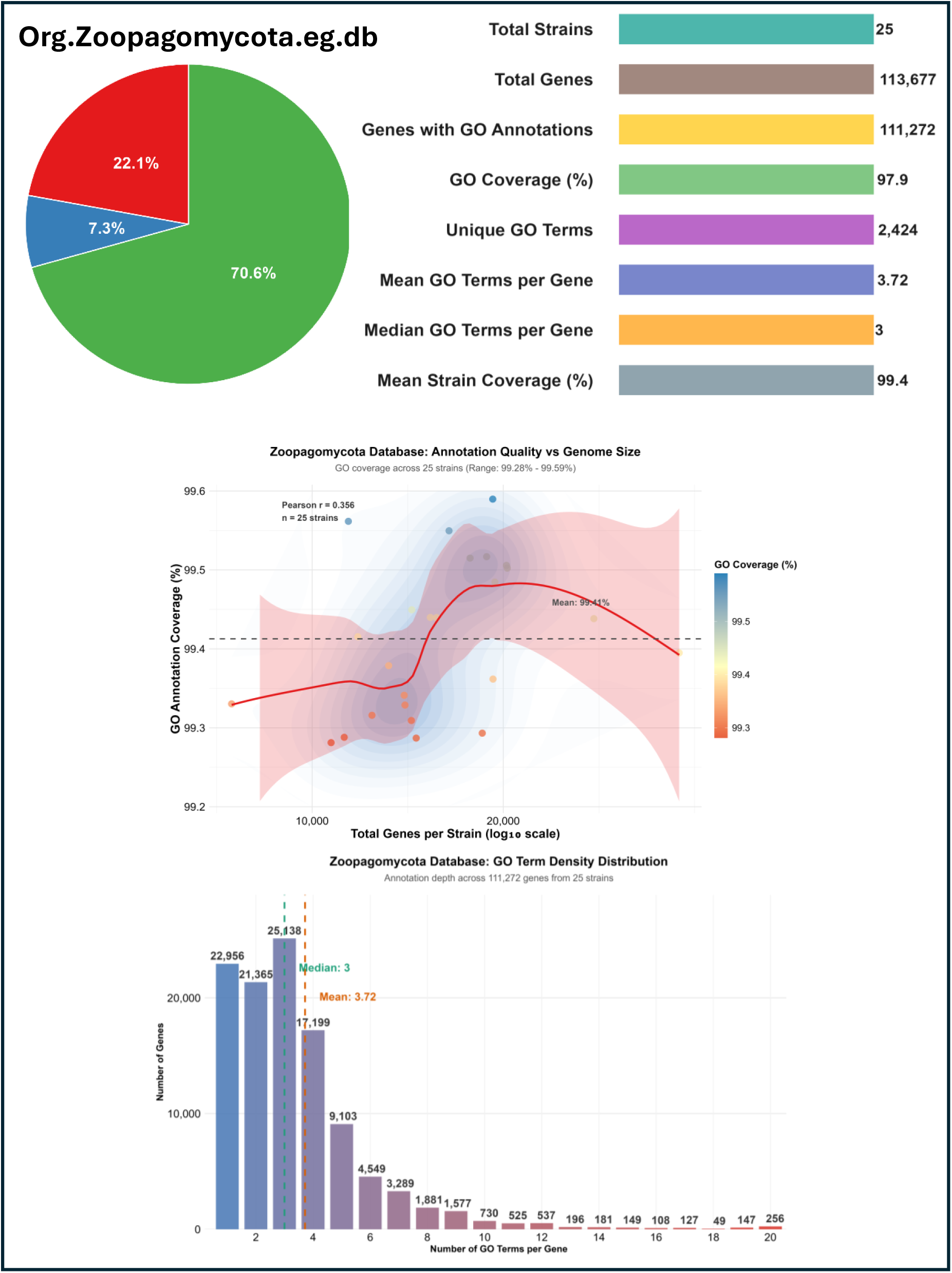
Gene Ontology Annotation Architecture of the Zoopagomycota Database. Comprehensive overview of Org.Zoopagomycota.eg.db representing 25 strains from early-diverging fungal lineages. **Top panel**: Database composition and GO annotation metrics. **Pie chart**: Distribution across Biological Process (red), Molecular Function (green), and Cellular Component (blue) categories. **Middle panel**: Scatter plot with LOESS curve showing genome size versus GO coverage relationship (Pearson r = 0.356), with strains color-coded by coverage percentage and dashed line indicating mean. **Bottom panel**: Histogram of GO term frequency distribution per gene across 111,272 annotated genes, showing characteristic right-skewed pattern with peak at 3 terms per gene.

**Figure 2:**
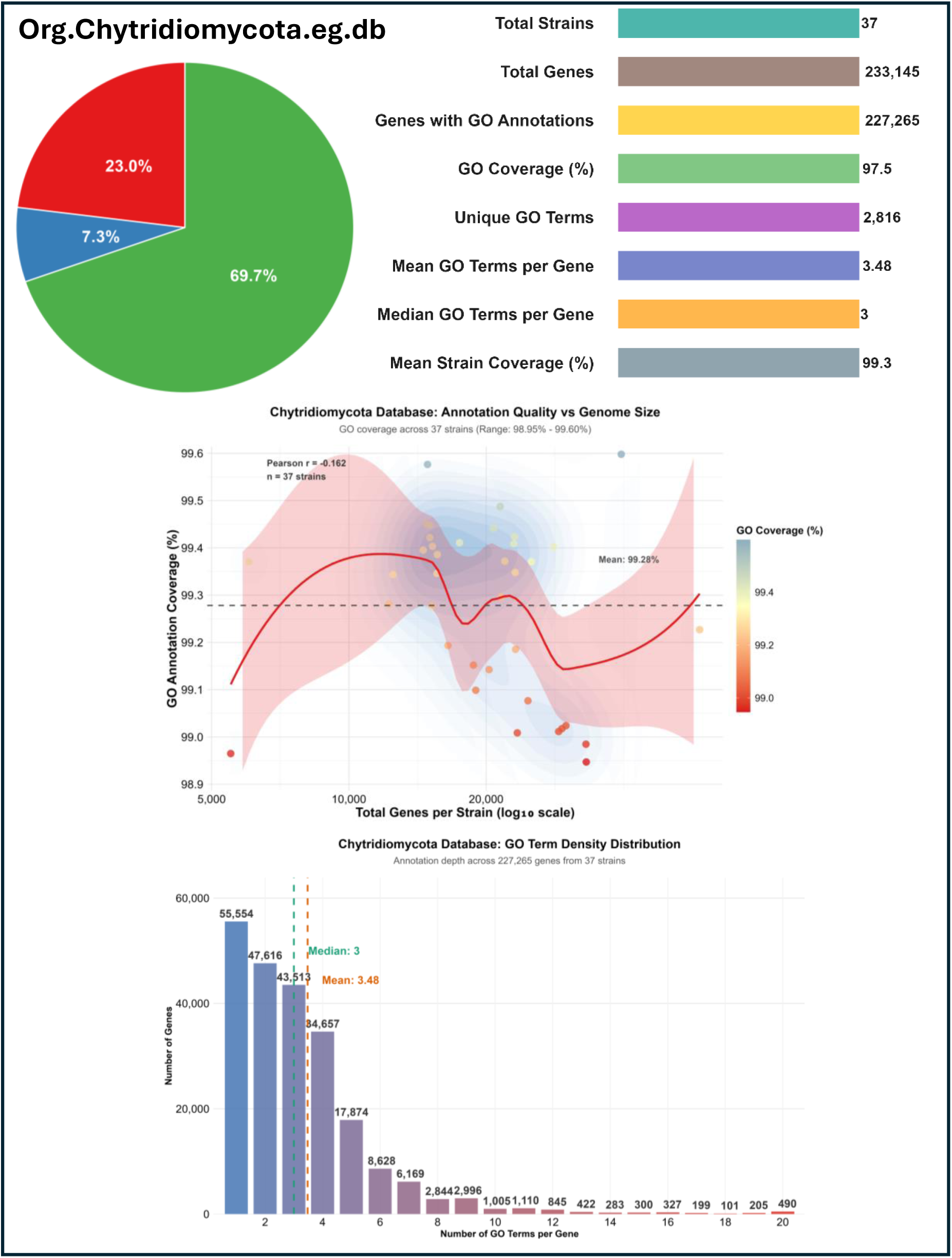
Functional Annotation Landscape of the Chytridiomycota Database. Functional annotation characteristics of Org.Chytridiomycota.eg.db representing flagellated aquatic fungi across 37 strains. **Top panel**: Database composition and annotation metrics. **Pie chart**: GO category distribution across the three major ontology branches. **Middle panel**: Scatter plot with LOESS curve showing genome size versus GO coverage relationship (Pearson r = −0.162), with strains color-coded by coverage percentage and dashed line indicating mean coverage. **Bottom panel**: Histogram of GO term frequency distribution per gene across 227,265 annotated genes, showing right-skewed pattern with modal value at 3 terms per gene.

**Figure 3:**
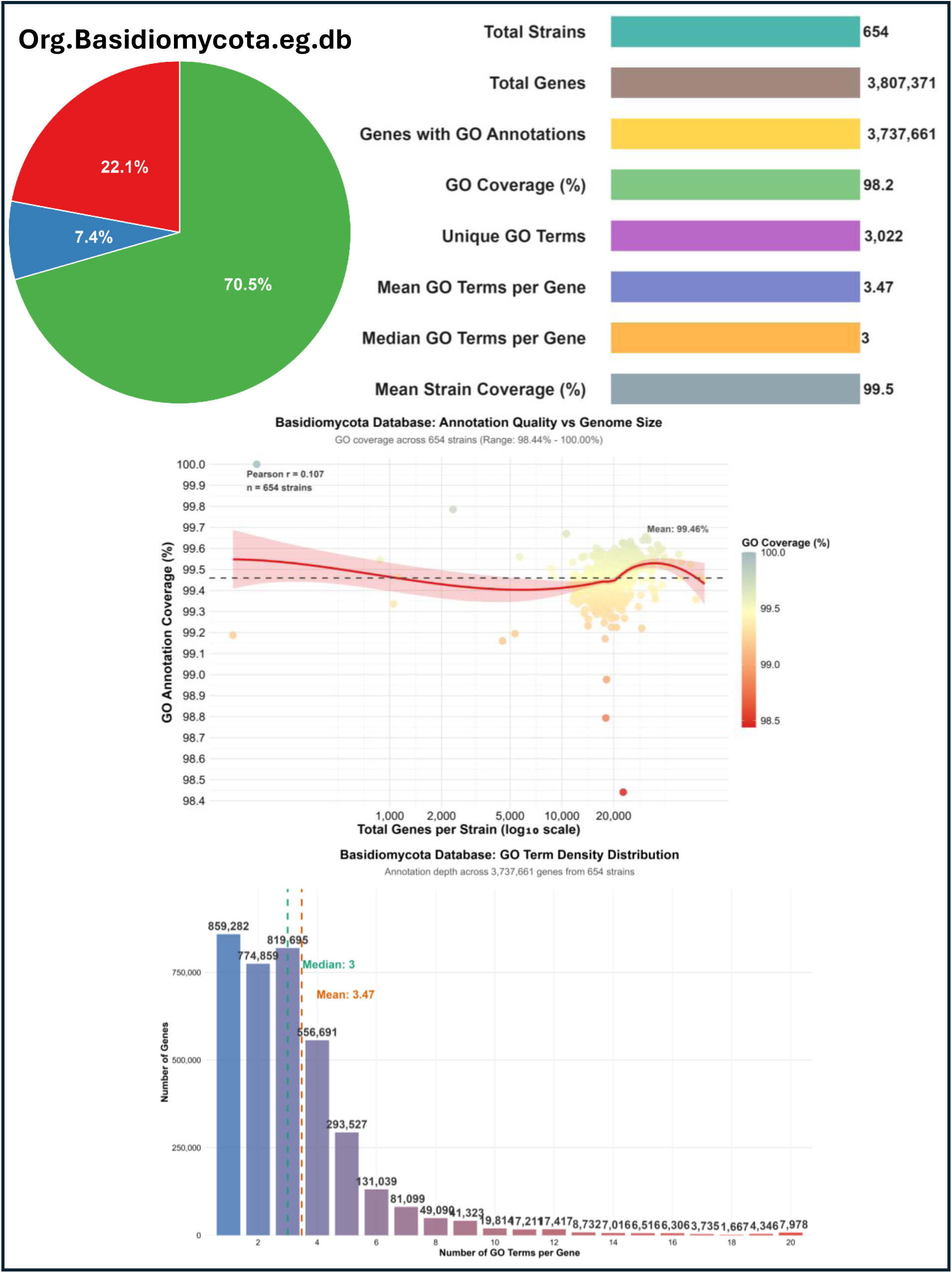
Mucoromycota Database Annotation Depth and Coverage Patterns. Functional annotation profile of Org.Mucoromycota.eg.db encompassing 172 strains from this ecologically diverse early-diverging lineage. **Top panel**: Database composition and GO annotation metrics. **Pie chart**: Distribution across Biological Process (red), Molecular Function(green), and Cellular Component(blue) categories. **Middle panel**: Scatter plot with LOESS curve showing genome size versus GO coverage relationship (Pearson r = −0.222), with strains color-coded by coverage percentage and dashed line indicating mean. **Bottom panel**: Histogram of GO term frequency distribution per gene across 974,709 annotated genes, showing characteristic right-skewed pattern with modal peak in the 1-3 term range.

**Figure 4:**
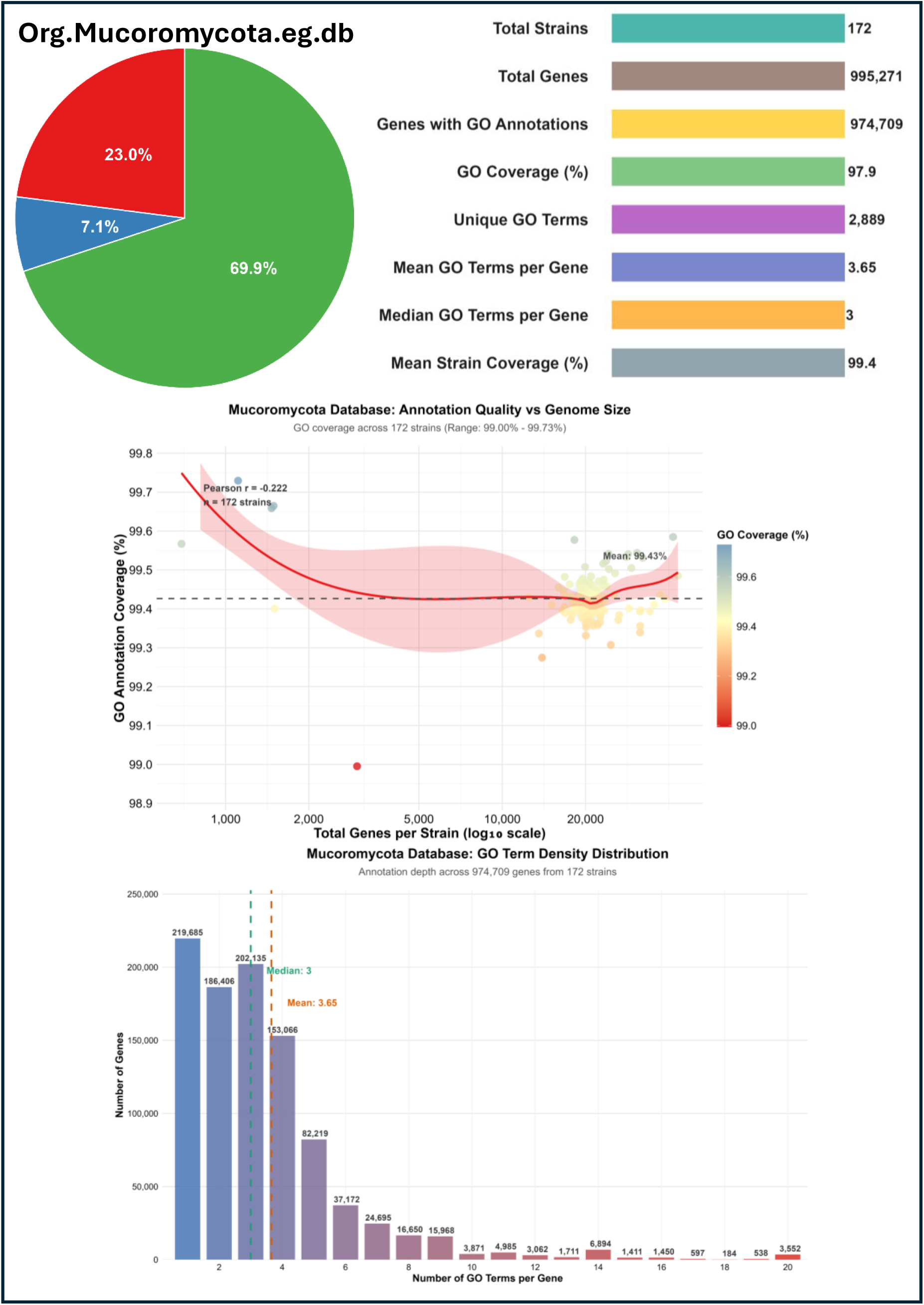
Basidiomycota Database Structure and Annotation Consistency. Functional annotation architecture of Org.Basidiomycota.eg.db representing 654 strains from this ecologically diverse phylum encompassing major decomposers, mycorrhizal symbionts, and plant pathogens. **Top panel**: Database composition and GO annotation metrics. **Pie chart**: GO category distribution across the three ontology branches: Biological Process (red), Molecular function (green) and Cellular components(blue). **Middle panel**: Scatter plot with LOESS curve showing genome size versus GO coverage relationship across 654 strains (Pearson r = 0.107), with strains color-coded by coverage percentage and dashed line indicating mean. **Bottom panel**: Histogram of GO term frequency distribution per gene across 3,737,661 annotated genes, showing right-skewed pattern with modal class at 3 terms per gene.

**Figure 5:**
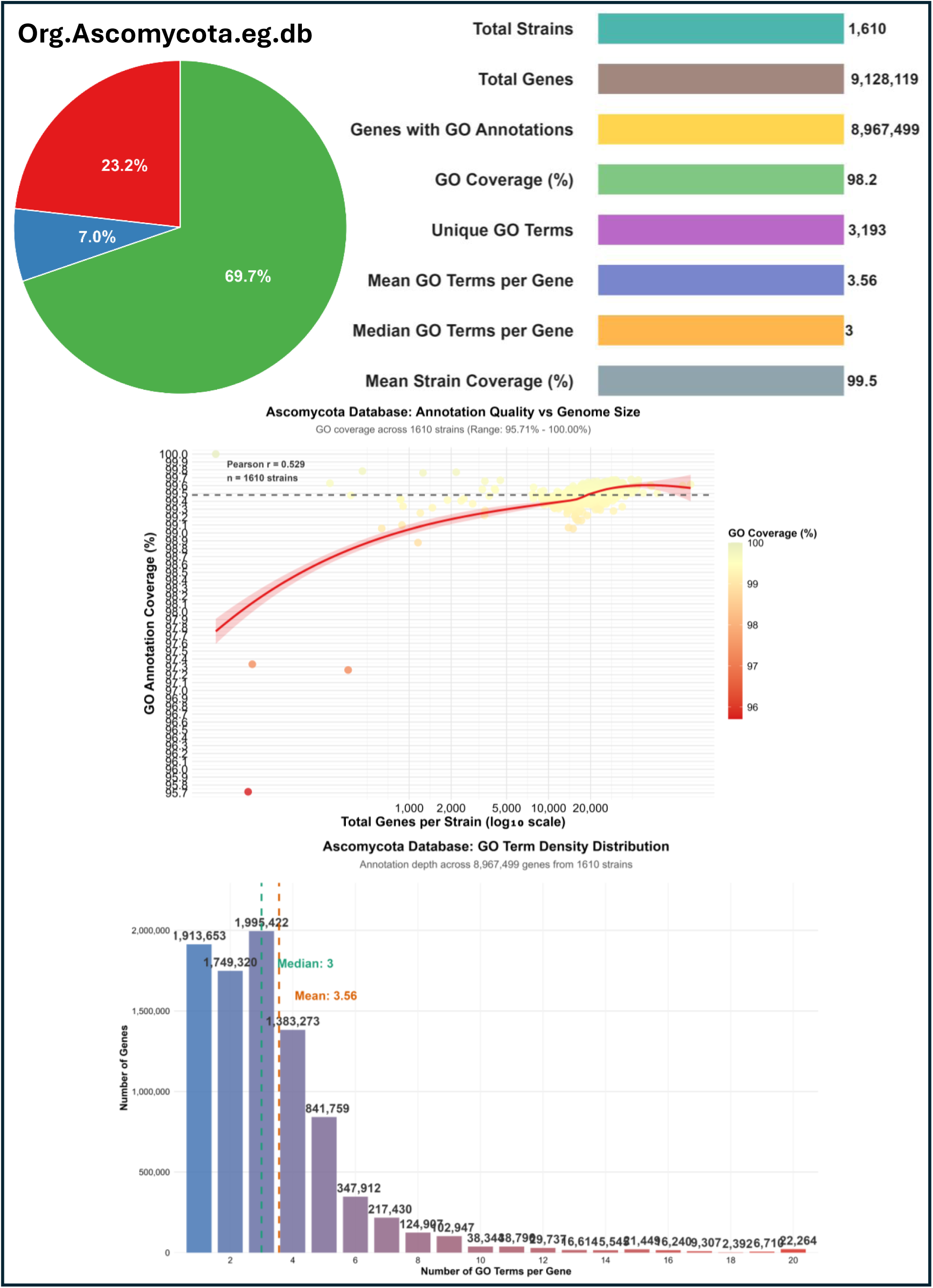
Ascomycota Database Scale and Annotation Coverage Dynamics. Comprehensive functional annotation landscape of Org.Ascomycota.eg.db representing 1,610 strains from this species-rich phylum including model organisms, plant pathogens, and industrial species. **Top panel**: Database composition and GO annotation metrics showing the largest strain collection among all databases. **Pie chart**: GO category distribution across ontology branches Biological Process (red), Molecular function (green) and Cellular components(blue). **Middle panel**: Scatter plot with LOESS curve showing genome size versus GO coverage relationship (Pearson r = 0.529), with strains color-coded by coverage percentage and dashed line indicating mean. The positive correlation suggests larger ascomycete genomes achieve slightly higher annotation completeness. **Bottom panel**: Histogram of GO term frequency distribution per gene across 8,967,499 annotated genes, demonstrating characteristic right-skewed pattern with modal value at 3 terms per gene despite massive database scale.

