## Supplementary Material for "Comprehensive OrgDb Packages for Fungal Comparative Genomics: MycoCosm-Derived Standardized GO and InterPro Annotations Across Five Major Phyla"

**Case Study 1**: Phanaerochaete chrysosporium and Postia placenta gene expression in ball-milled aspen or ball-milled pine medium [GSE29659]

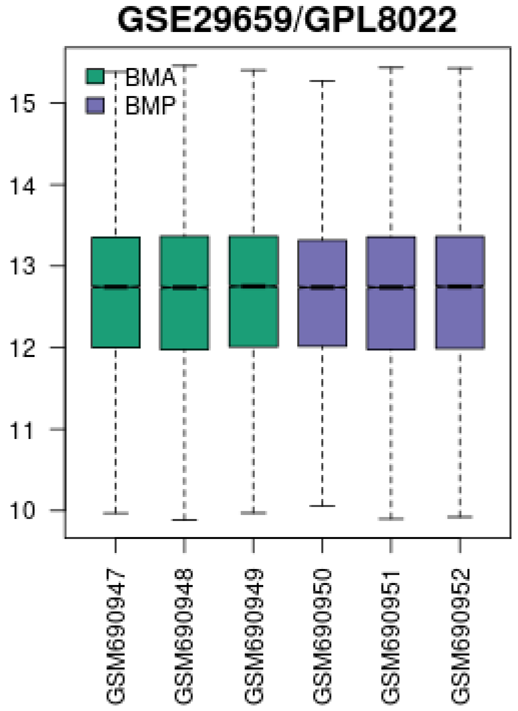


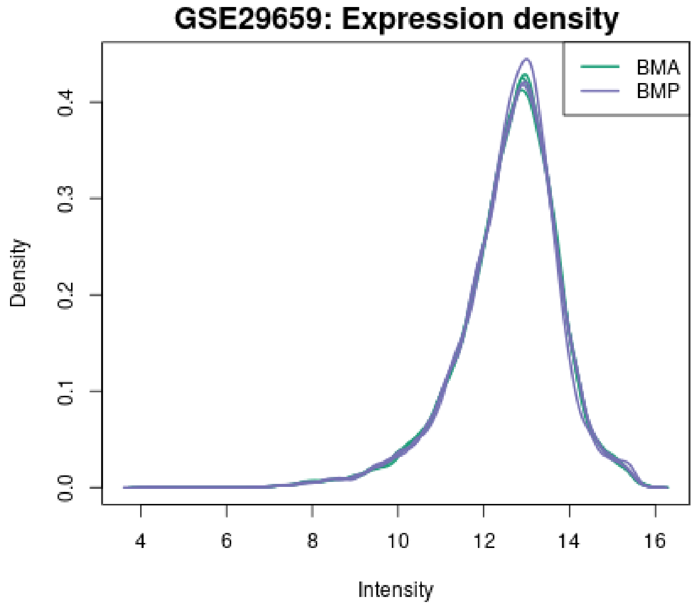


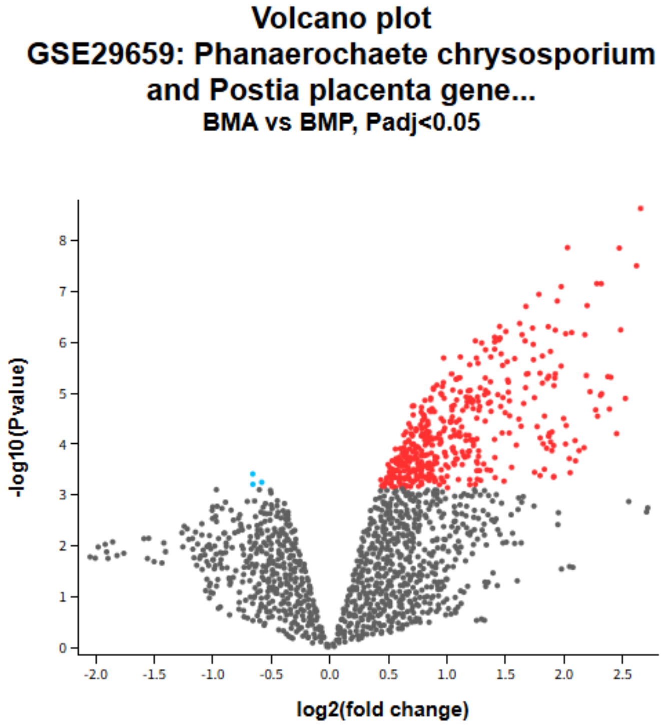

Steps to perform Gene Ontology Enrichment using the Differentially Expressed Genes obtained from the comparison of Ball Milled Aspen [BMA] (vs) Ball Milled Pine [BMP]- GSE29659

**Step-1**: Install the "org.Basidiomycota.eg.db" using the link

```
>install.packages("devtools")

>devtools::install_github("ayistak89/Org.Basidiomycota.eg.db")

>library (org.Basidiomycota.eg.db)
```

**Step-2**: After loading the package retrieve all the genes present in the database

```
>all_genes <- keys (org.Basidiomycota.eg.db, "GID")
```

**Step-3:** As our present study is focussed on *Phanerochaete chrysosporium* fungal strain we will be retrieving all the genes of *Phanerochaete chrysosporium* "Phchr2" using:

```
>phchr2_genes <- all_genes[grepl("^Phchr2", all_genes)]

> length(phchr2_genes)
```

**Step-4:** Now we have loaded the background genes (`phchr2_genes`) we will be importing the differentially expressed genes obtained from the comparison of BMA (vs) BMP of *P. chrysosporium* cultures.

**Step-5:** Import the DEGs using read.csv ("/path/to_the/location/GSE29659_DEGs.csv", sep = ",") [or]

```
GSE29659_DEGs <- unique(c("Phchr2_2226","Phchr2_4052","Phchr2_2226",

"Phchr2_2226","Phchr2_10276","Phchr2_10276","Phchr2_4052","Phchr2_10551","Phc
hr2_4052","Phchr2_10276","Phchr2_138479","Phchr2_138479","Phchr2_126","Phchr2
_6458","Phchr2_138479","Phchr2_5517","Phchr2_8996","Phchr2_138266","Phchr2_40
899","Phchr2_40899","Phchr2_138266","Phchr2_40899","Phchr2_3686","Phchr2_5571
","Phchr2_140079","Phchr2_126","Phchr2_121582","Phchr2_8759","Phchr2_1903","P
hchr2_130517","Phchr2_10551","Phchr2_8466","Phchr2_8466","Phchr2_131217","Phc
hr2_134001","Phchr2_3518","Phchr2_5571","Phchr2_7398","Phchr2_138021","Phchr2
```

_121582","Phchr2_138266","Phchr2_4361","Phchr2_140079","Phchr2_1903","Phchr2_
130517","Phchr2_126","Phchr2_8996","Phchr2_6458","Phchr2_10551","Phchr2_13144
0","Phchr2_7398","Phchr2_3518","Phchr2_2450","Phchr2_131217","Phchr2_5517","P
hchr2_5517","Phchr2_6153","Phchr2_140079","Phchr2_4550","Phchr2_131217","Phch
r2_6458","Phchr2_5249","Phchr2_3328","Phchr2_138350","Phchr2_41650","Phchr2_1
34001","Phchr2_1165","Phchr2_3518","Phchr2_133585","Phchr2_131440","Phchr2_11
39","Phchr2_8759","Phchr2_5873","Phchr2_5145","Phchr2_138021","Phchr2_3328","
Phchr2_139063","Phchr2_41650","Phchr2_5873","Phchr2_2450","Phchr2_4468","Phch
r2_1903","Phchr2_138350","Phchr2_138021","Phchr2_140501","Phchr2_2710","Phchr
2_134001","Phchr2_41650","Phchr2_2450","Phchr2_134615","Phchr2_127396","Phchr
2_5571","Phchr2_131440","Phchr2_121582","Phchr2_5145","Phchr2_1165","Phchr2_1
21077","Phchr2_126191","Phchr2_3328","Phchr2_2544","Phchr2_7054","Phchr2_1744
","Phchr2_8466","Phchr2_10833","Phchr2_2544","Phchr2_138350","Phchr2_127396",
"Phchr2_3686","Phchr2_4361","Phchr2_4550","Phchr2_3854","Phchr2_8916","Phchr2
_6153","Phchr2_1744","Phchr2_6153","Phchr2_2925","Phchr2_1696","Phchr2_130517
","Phchr2_128442","Phchr2_121077","Phchr2_140501","Phchr2_612","Phchr2_10833"
,"Phchr2_134492","Phchr2_126191","Phchr2_8311","Phchr2_133585","Phchr2_7852",
"Phchr2_5787","Phchr2_6482","Phchr2_10009","Phchr2_2991","Phchr2_7054","Phchr
2_6482","Phchr2_8996","Phchr2_140501","Phchr2_612","Phchr2_126191","Phchr2_17
44","Phchr2_10607","Phchr2_35408","Phchr2_8759","Phchr2_3713","Phchr2_5145","
Phchr2_8470","Phchr2_612","Phchr2_10833","Phchr2_7398","Phchr2_6893","Phchr2_
127920","Phchr2_1165","Phchr2_5607","Phchr2_1696","Phchr2_134145","Phchr2_530
6","Phchr2_10607","Phchr2_138589","Phchr2_4361","Phchr2_127920","Phchr2_13427
7","Phchr2_4913","Phchr2_4550","Phchr2_5607","Phchr2_600","Phchr2_4468","Phch
r2_138825","Phchr2_8570","Phchr2_133585","Phchr2_3805","Phchr2_121155","Phchr
2_5607","Phchr2_139063","Phchr2_6482","Phchr2_127396","Phchr2_138589","Phchr2
_10202","Phchr2_4796","Phchr2_140807","Phchr2_121077","Phchr2_1139","Phchr2_1
38345","Phchr2_134145","Phchr2_2925","Phchr2_8916","Phchr2_133992","Phchr2_16
96","Phchr2_139063","Phchr2_5873","Phchr2_134492","Phchr2_3761","Phchr2_5365"
,"Phchr2_2936","Phchr2_2544","Phchr2_4796","Phchr2_1414","Phchr2_137747","Phc
hr2_7852","Phchr2_127920","Phchr2_8470","Phchr2_843","Phchr2_3805","Phchr2_27
10","Phchr2_1139","Phchr2_968","Phchr2_3761","Phchr2_3686","Phchr2_128442","P
hchr2_8935","Phchr2_2925","Phchr2_990","Phchr2_8254","Phchr2_5306","Phchr2_52
49","Phchr2_137216","Phchr2_27652","Phchr2_10607","Phchr2_138345","Phchr2_524
9","Phchr2_5306","Phchr2_121155","Phchr2_134145","Phchr2_4796","Phchr2_122292
","Phchr2_124963","Phchr2_8688","Phchr2_3761","Phchr2_3171","Phchr2_34295","P
hchr2_138589","Phchr2_4913","Phchr2_134615","Phchr2_137747","Phchr2_44722","P
hchr2_10375","Phchr2_3713","Phchr2_2991","Phchr2_27652","Phchr2_124963","Phch
r2_4307","Phchr2_3713","Phchr2_2441","Phchr2_136839","Phchr2_10009","Phchr2_4
307","Phchr2_1414","Phchr2_2035","Phchr2_3805","Phchr2_147","Phchr2_124963","
Phchr2_134277","Phchr2_6766","Phchr2_6766","Phchr2_121806","Phchr2_5319","Phc
hr2_8935","Phchr2_9006","Phchr2_2035","Phchr2_8570","Phchr2_3630","Phchr2_847
0","Phchr2_34295","Phchr2_3854","Phchr2_32547","Phchr2_35408","Phchr2_10189",
"Phchr2_147","Phchr2_8935","Phchr2_134277","Phchr2_134615","Phchr2_147","Phch
r2_134492","Phchr2_8311","Phchr2_136963","Phchr2_2722","Phchr2_7809","Phchr2_
8617","Phchr2_10009","Phchr2_38357","Phchr2_34295","Phchr2_137372","Phchr2_69
29","Phchr2_5787","Phchr2_137216","Phchr2_140807","Phchr2_2220","Phchr2_12702
9","Phchr2_5499","Phchr2_6211","Phchr2_2722","Phchr2_127717","Phchr2_6893","P
hchr2_122292","Phchr2_133992","Phchr2_133757","Phchr2_35","Phchr2_122324","Ph

```
chr2_10757","Phchr2_10375","Phchr2_7852","Phchr2_138749","Phchr2_5348","Phchr
2_3361","Phchr2_8570","Phchr2_137372","Phchr2_122292","Phchr2_2710","Phchr2_4
4722","Phchr2_4874","Phchr2_5348","Phchr2_2117","Phchr2_139612","Phchr2_2220"
,"Phchr2_1748","Phchr2_1450","Phchr2_121155","Phchr2_4468","Phchr2_133813","P
hchr2_6211","Phchr2_138710","Phchr2_843","Phchr2_132680","Phchr2_136963","Phc
hr2_990","Phchr2_138825","Phchr2_7029","Phchr2_6893","Phchr2_140323","Phchr2_
5319","Phchr2_5787","Phchr2_138749","Phchr2_7054","Phchr2_8712","Phchr2_5352"
,"Phchr2_27652","Phchr2_137747","Phchr2_6211","Phchr2_41616","Phchr2_8346","P
hchr2_4874","Phchr2_136963","Phchr2_628","Phchr2_135167","Phchr2_137372","Phc
hr2_4874","Phchr2_600","Phchr2_6766","Phchr2_136145","Phchr2_138710","Phchr2_
128442","Phchr2_129325","Phchr2_8052","Phchr2_1088","Phchr2_2441","Phchr2_299
1","Phchr2_137216","Phchr2_4307","Phchr2_5348","Phchr2_138838","Phchr2_7029",
"Phchr2_4672","Phchr2_2441","Phchr2_134114","Phchr2_121806","Phchr2_5126","Ph
chr2_35408","Phchr2_6997","Phchr2_122095","Phchr2_8688","Phchr2_8712","Phchr2
_133757","Phchr2_138710","Phchr2_138397","Phchr2_3854","Phchr2_133757","Phchr
2_25018","Phchr2_41330","Phchr2_8052","Phchr2_7548","Phchr2_7633","Phchr2_250
18","Phchr2_1450","Phchr2_138784","Phchr2_10189","Phchr2_135167","Phchr2_7809
","Phchr2_127029","Phchr2_8916","Phchr2_6997","Phchr2_32547","Phchr2_41330","
Phchr2_10202","Phchr2_8311","Phchr2_26890","Phchr2_2035","Phchr2_2220","Phchr
2_8346","Phchr2_968","Phchr2_5054","Phchr2_1308","Phchr2_127029","Phchr2_968"
,"Phchr2_5774","Phchr2_139777","Phchr2_122324","Phchr2_7633","Phchr2_2569","P
hchr2_8688","Phchr2_4180","Phchr2_8604","Phchr2_136839","Phchr2_129325","Phch
r2_133992","Phchr2_1088","Phchr2_133020","Phchr2_3361","Phchr2_5024","Phchr2_
126443","Phchr2_37620","Phchr2_25018","Phchr2_134556","Phchr2_2011","Phchr2_3
2547","Phchr2_140491","Phchr2_600","Phchr2_122324","Phchr2_5747","Phchr2_1293
25","Phchr2_2936","Phchr2_133020","Phchr2_6579","Phchr2_8346","Phchr2_1231","
Phchr2_136145","Phchr2_133813","Phchr2_5155","Phchr2_35","Phchr2_1450","Phchr
2_136839"))
```

**Step-6:** Using the DEGs obtained to perform the gene ontology of biological processes:

```
> GSE29659_DEGs_BP <- enrichGO_Basidiomycota(gene = GSE29659_DEGs,
                      universe = phchr2_genes, ont = "BP",
                      pvalueCutoff = 0.05, qvalueCutoff = 0.05,
                      minGSSize = 5)

> p1 <- barplot(GSE29659_DEGs_BP, showCategory = 10)

> p <- pairwise_termsim(GSE29659_DEGs_BP)

> p2 <-cnetplot(p, showCategory = 10, colorEdge = TRUE, circular =
FALSE, node_label = "all")
```

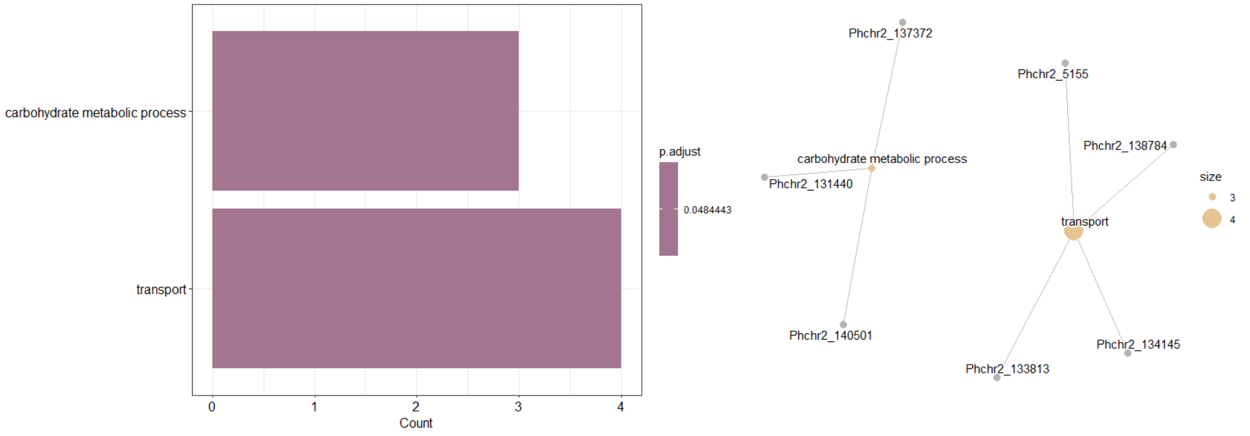


**Step-7:** Using the DEGs obtained to perform the gene ontology of Molecular Function:

```
> GSE29659_DEGs_MF <- enrichGO_Basidiomycota(gene = GSE29659_DEGs,
                    universe = phchr2_genes, ont = "MF",
                    pvalueCutoff = 0.05, qvalueCutoff = 0.05,
                    minGSSize = 5)

> p1 <- barplot(GSE29659_DEGs_MF, showCategory = 10)

> p <- pairwise_termsim(GSE29659_DEGs_MF)

> p2 <-cnetplot(p, showCategory = 10, colorEdge = TRUE, circular =
FALSE, node_label = "all")
```

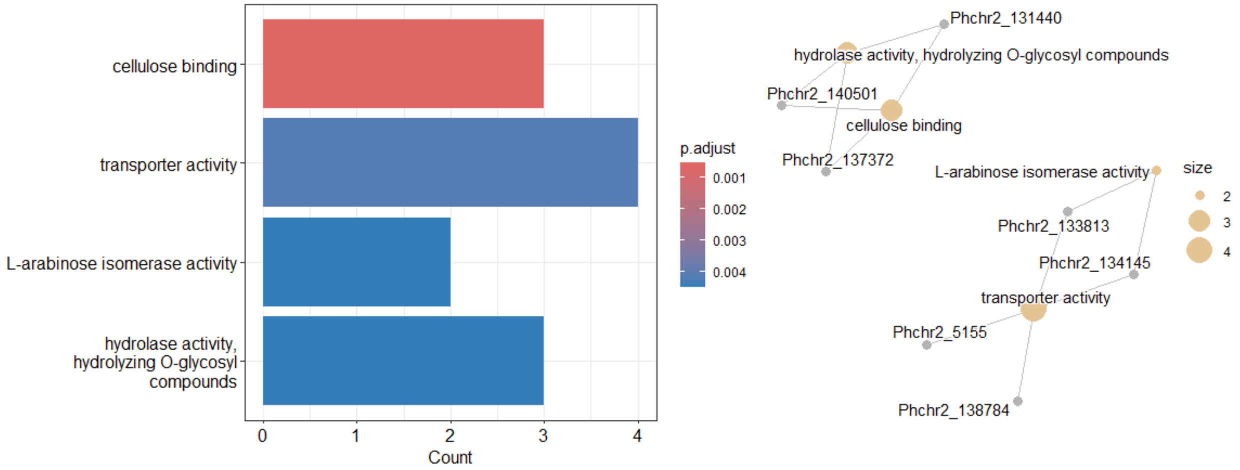

**Step-8:** Using the DEGs obtained to perform the gene ontology of Cellular component:

```
> GSE29659_DEGs_CC <- enrichGO_Basidiomycota(gene = GSE29659_DEGs,
                      universe = phchr2_genes, ont = "CC",
                      pvalueCutoff = 0.05, qvalueCutoff = 0.05,
                      minGSSize = 5)

> p1 <- barplot(GSE29659_DEGs_CC, showCategory = 10)

> p <- pairwise_termsim(GSE29659_DEGs_CC)

> p2 <-cnetplot(p, showCategory = 10, colorEdge = TRUE, circular =
FALSE, node_label = "all")
```

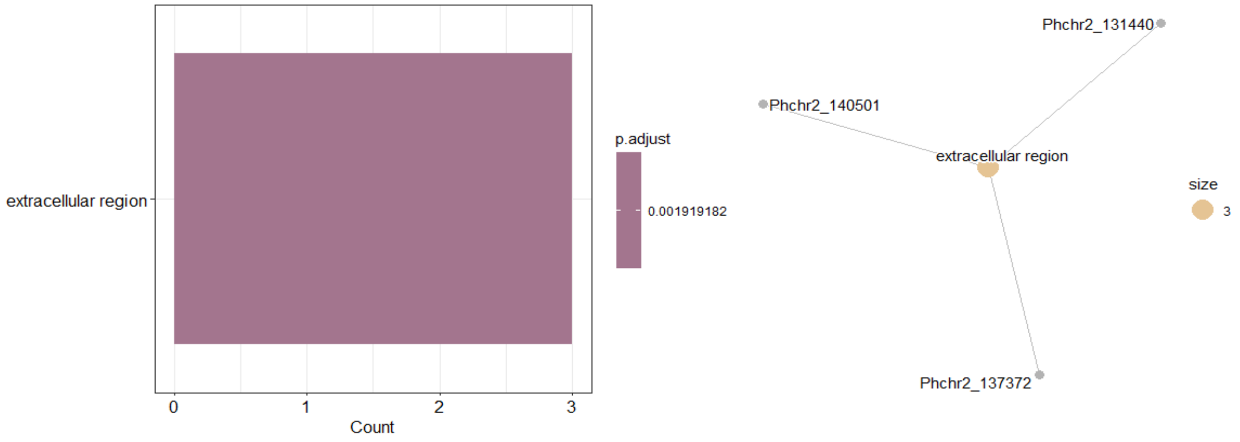

**<u>Case Study</u> 2: Analyzing target and comparison strains**

```
> target strain <- "Neosp1"

Option-A :

> comparison_strains <- c("Neolan1", "Neocon1", "Orpsp1", "PirE2",
"Pirfi3", "Anasp1", "Piromy")


> unique_results <- analyze_unique_go_terms(
                     target_strain = target_strain,
                     comparison_strains = comparison_strains,
                     sample_size = 500,
                     pvalue_cutoff = 0.05,
                     uniqueness_threshold = 0.1)


> barplot_bp <- create_custom_barplot(
  unique_results$unique_terms$BP,
  title = paste("Unique Biological Processes in", target_strain),
  analysis_type = "unique",
  top_n = 15)
```

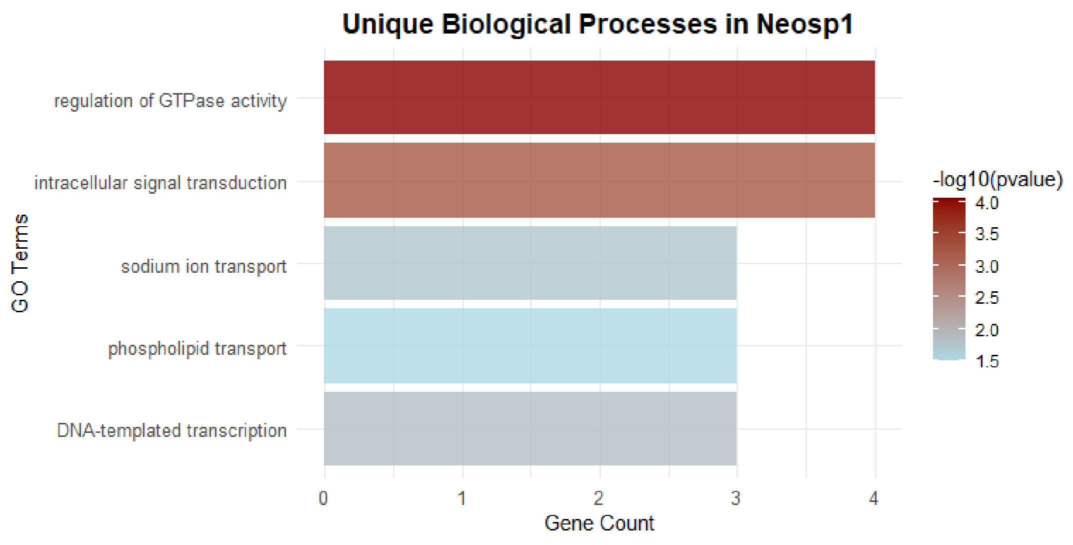

```
> barplot_MF <- create_custom_barplot(
  unique_results$unique_terms$MF,
  title = paste("Unique Molecular Function in", target_strain),
  analysis_type = "unique",
  top_n = 15)

> print(barplot_MF)
```

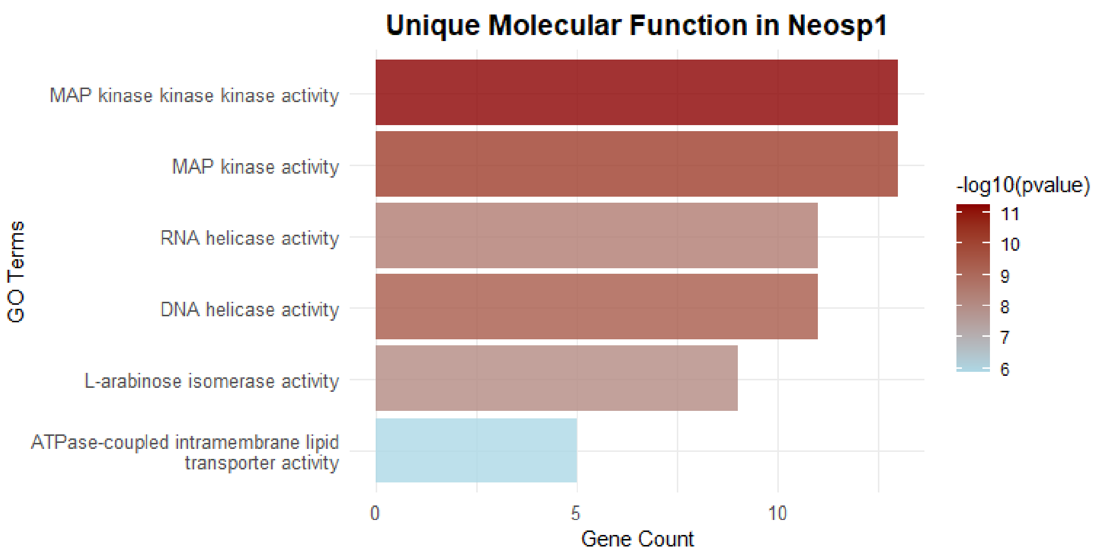


```
> barplot_CC <- create_custom_barplot(
  unique_results$unique_terms$CC,
  title = paste("Unique Cellular Components in", target_strain),
  analysis_type = "unique",
  top_n = 15)

> print(barplot_CC)
```

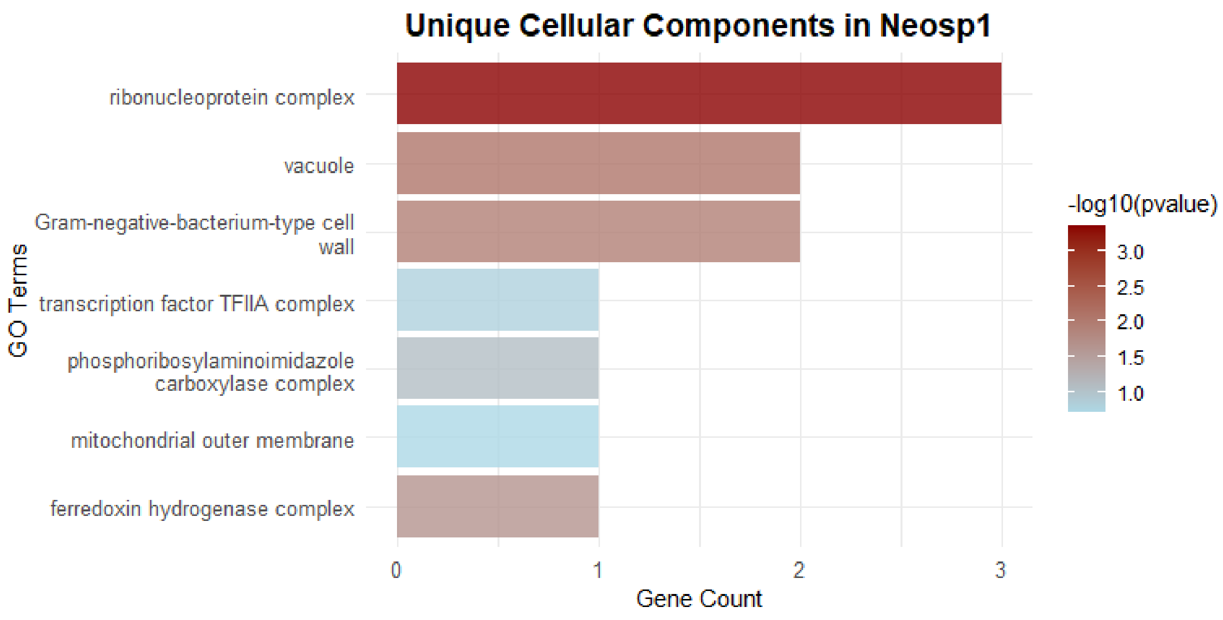


**<u>CASE Study -3:</u> Conserved Gene Ontology enrichment in selected strains**

```
> strains <- unique(available_strains$strain)

> strains

[1] "Neosp1"   "NeoGfMa1" "NeoWI3"   "Ganpr1"   "Neolan1"
"Neocon1"  "Orpsp1"   "Pecora1"  "Rhihy1"   "Clarep1"  "Hyacur1"
"Chyhya1" "Piromy"   "Chytri1"  "PirE2"    "Clapol1"  "Obemuc1"
"Caecom1"  "Pirfi3"   "Entlut1"  "Glopol1"  "Spipu1"   "Chylag1"
"Polagg1" "Fimjon1"  "Powhir1"  "Triarc1"  "Gervar1"  "Enthel1"
"Gaesem1"  "Gorhay1"  "Batde5"   "Blyhe1"   "Caupr1"   "Hompol1"
"Caupr" "Anasp1"


> conserved_results <- analyze_conserved_go_terms(
  selected_strains = strains,
  min_strains = 10,
  sample_size = 500,
  pvalue_cutoff = 0.05,
  max_strains = 37)
```

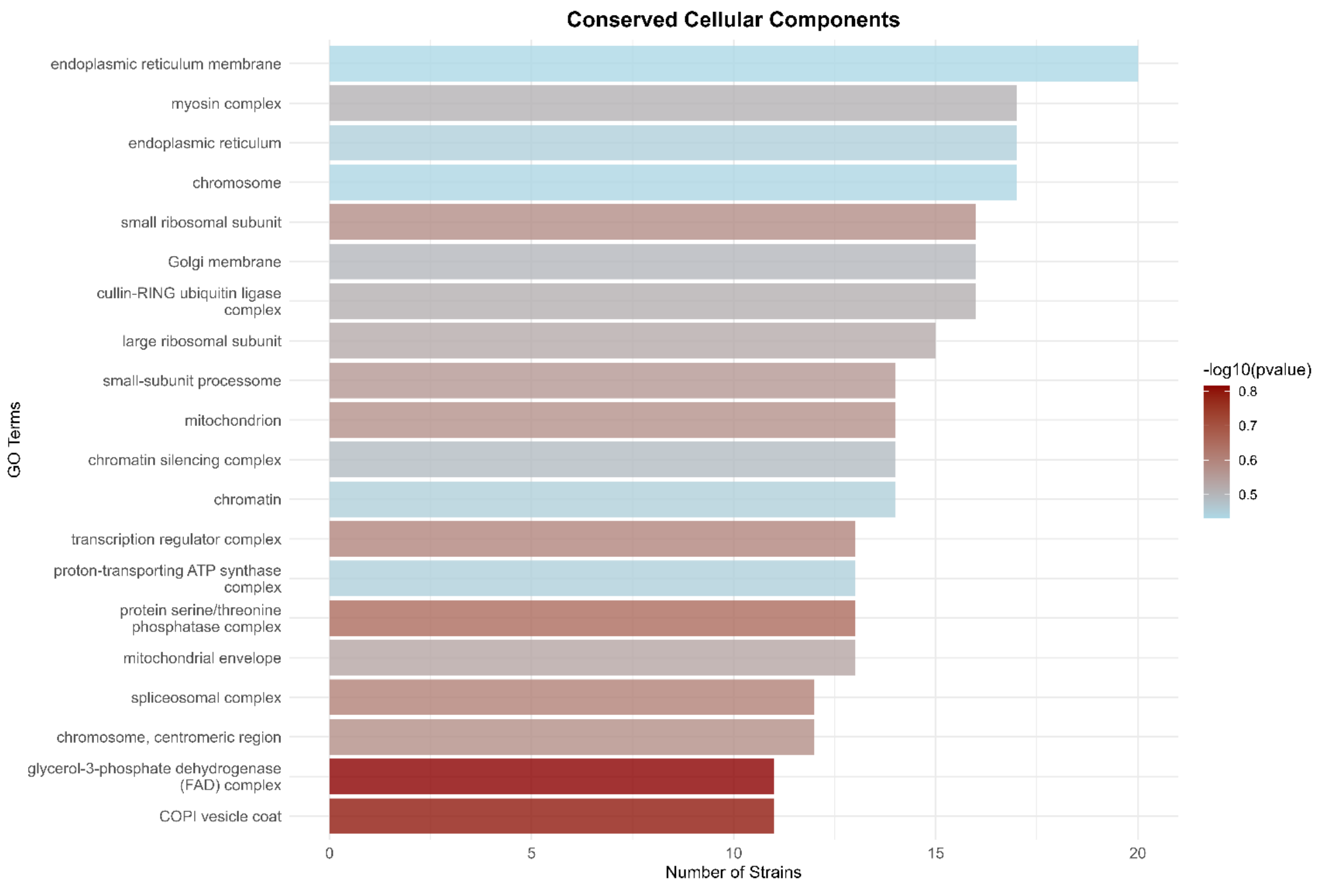


The function "`analyze_conserved_go_terms`" also performs the complete individual gene ontology enrichment in each individual fungal strain and then consolidates to analyze the conserved gene ontology enrichment among all the selected fungal strains. So this information is stored in:

```
>Neosp1_bp <- conserved_results$strain_results$BP$Neosp1

> Neosp1_bp <- create_custom_barplot(
  neosp1_bp,
  title = "Biological Processes in Neosp1",
  analysis_type = "unique",
  top_n = 20)

> print(Neosp1_bp)
```

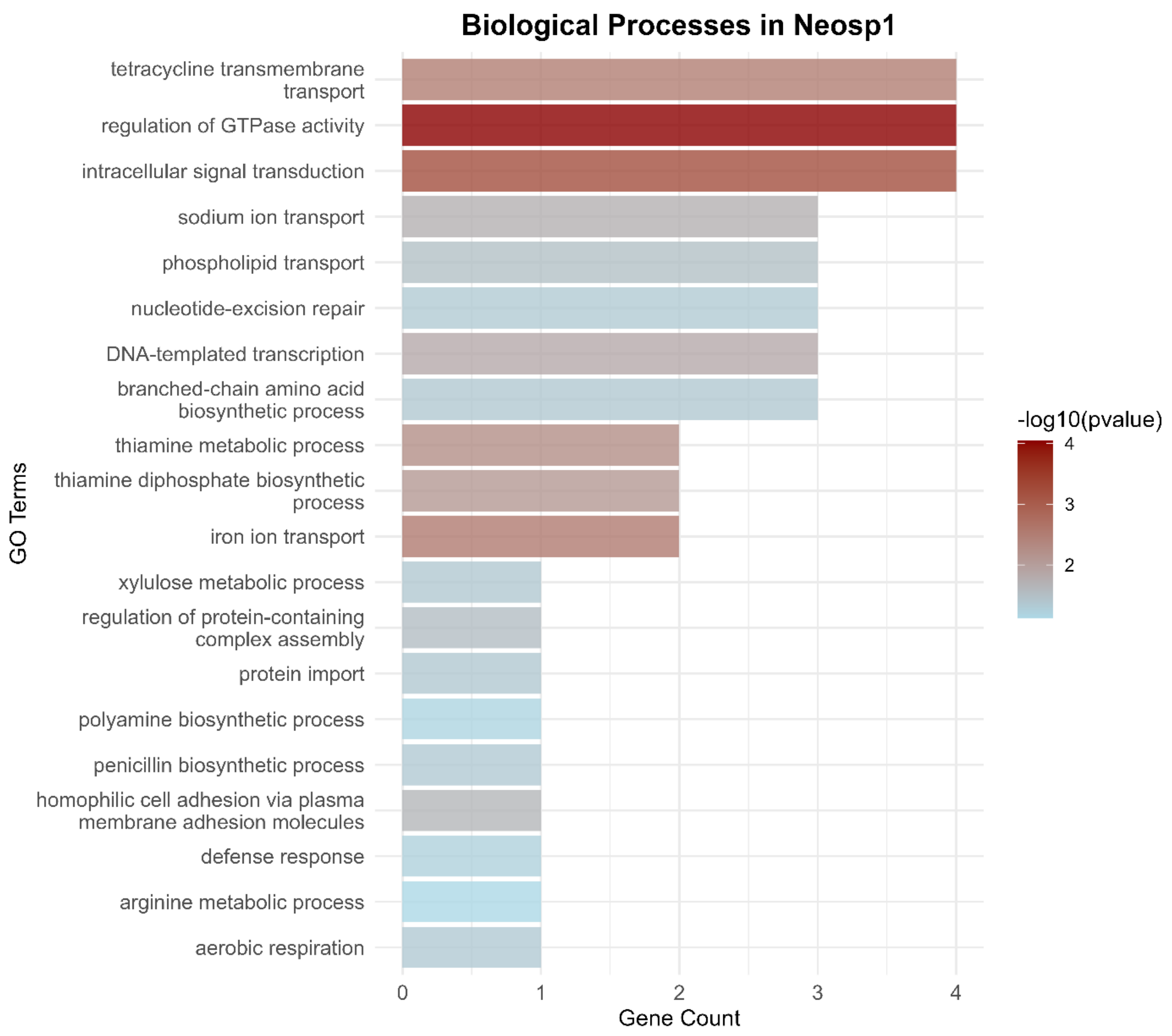


```
>Neosp1_CC <- conserved_results$strain_results$CC$Neosp1

> Neosp1_CC <- create_custom_barplot(
  Neosp1_CC,
  title = "Cellular Components in Neosp1",
  analysis_type = "unique",
  top_n = 20)

> print(Neosp1_CC)
```

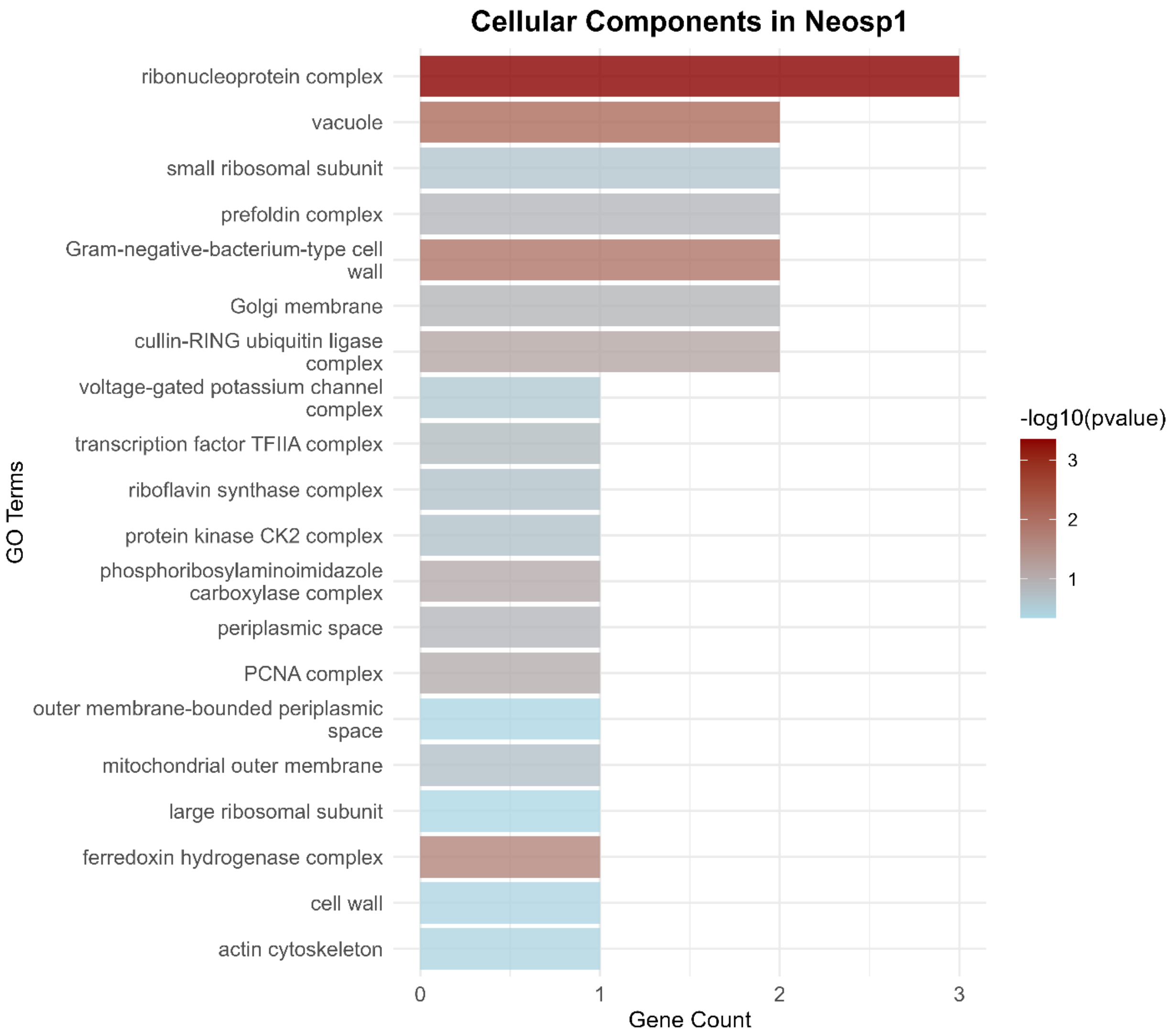


```
> Neosp1_MF <- conserved_results$strain_results$MF$Neosp1

> Neosp1_MF <- create_custom_barplot(
  Neosp1_MF,
  title = "Molecular Function in Neosp1",
  analysis_type = "unique",
  top_n = 20)

> print(Neosp1_MF)
```

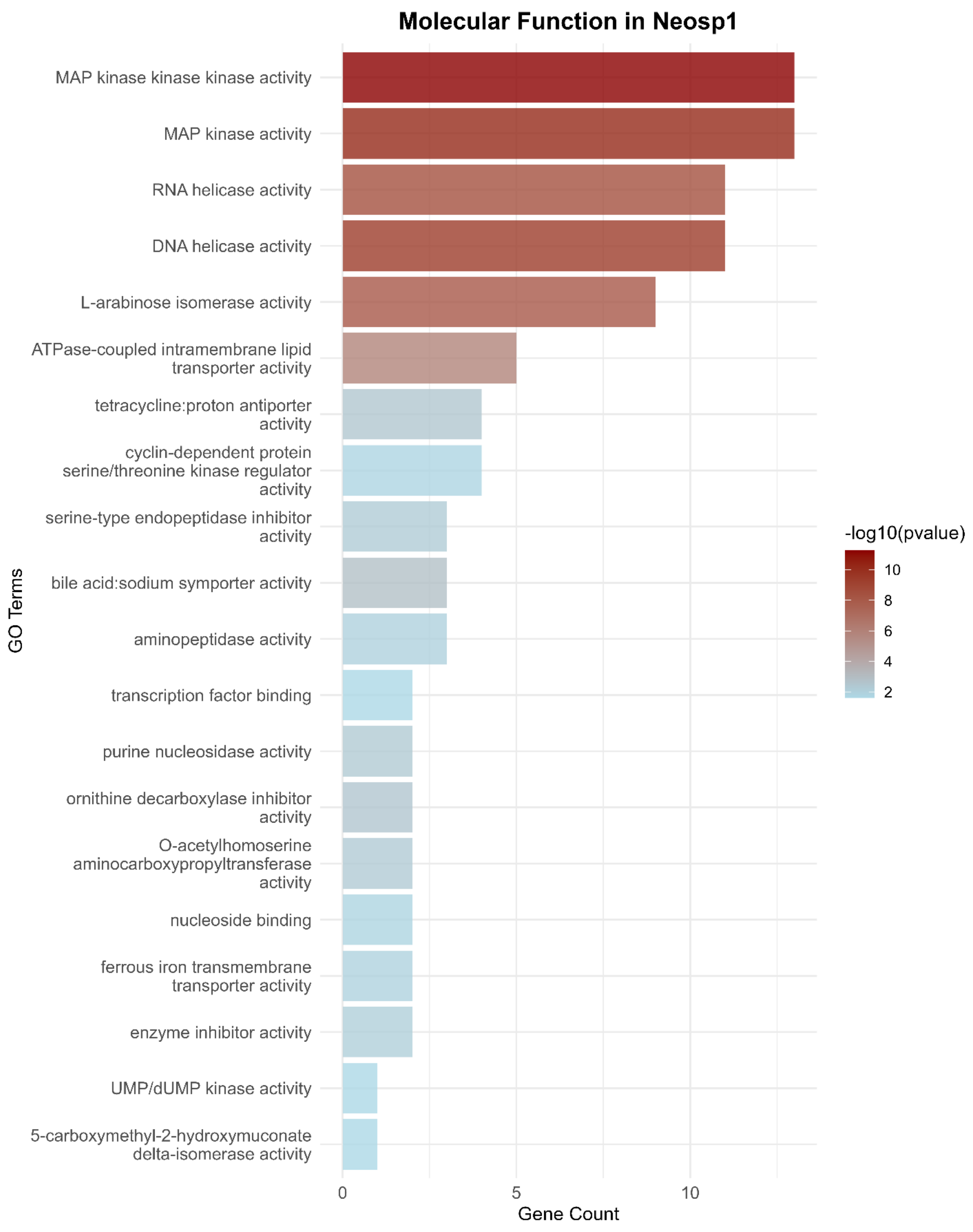
Molecular Function in Neosp1
MAP kinase kinase kinase activity
MAP kinase activity
RNA helicase activity
DNA helicase activity
L-arabinose isomerase activity
ATPase-coupled intramembrane lipid transporter activity
tetracycline:proton antiporter activity
cyclin-dependent protein serine/threonine kinase regulator activity
serine-type endopeptidase inhibitor activity
bile acid:sodium symporter activity
aminopeptidase activity
transcription factor binding
purine nucleosidase activity
ornithine decarboxylase inhibitor activity
O-acetylhomoserine aminocarboxypropyltransferase activity
nucleoside binding
ferrous iron transmembrane transporter activity
enzyme inhibitor activity
UMP/dUMP kinase activity
5-carboxymethyl-2-hydroxymuconate delta-isomerase activity
GO Terms
0
5
10
Gene Count
-log10(pvalue)
10
8
6
4
2
